# Distinct and Common Neural Coding of Semantic and Non-semantic Control Demands

**DOI:** 10.1101/2020.11.16.384883

**Authors:** Zhiyao Gao, Li Zheng, Rocco Chiou, André Gouws, Katya Krieger-Redwood, Xiuyi Wang, Dominika Varga, Matthew A. Lambon Ralph, Jonathan Smallwood, Elizabeth Jefferies

## Abstract

The flexible retrieval of knowledge is critical in everyday situations involving problem solving, reasoning and social interaction. Current theories emphasise the importance of a left-lateralised semantic control network (SCN) in supporting flexible semantic behaviour, while a bilateral multiple-demand network (MDN) is implicated in executive functions across domains. No study, however, has examined whether semantic and non-semantic demands are reflected in a common neural code within regions specifically implicated in semantic control. Using functional MRI and univariate parametric modulation analysis as well as multivariate pattern analysis, we found that semantic and non-semantic demands gave rise to both similar and distinct neural responses across control-related networks. Though activity patterns in SCN and MDN could decode the difficulty of both semantic and verbal working memory decisions, there was no shared common neural coding of cognitive demands in SCN regions. In contrast, regions in MDN showed common patterns across manipulations of semantic and working memory control demands, with successful cross-classification of difficulty across tasks. Therefore, SCN and MDN can be dissociated according to the information they maintain about cognitive demands.

## Introduction

Our semantic knowledge encompasses disparate features and associations for any given concept (e.g., APPLE can go with PIE but also HORSE). While this allows us to understand the significance of diverse experiences, it raises the question of how we generate coherent patterns of semantic retrieval that diverge from strong associations in the semantic store. The controlled semantic cognition framework suggests that a distributed neural network manipulates activation within the semantic representational system to generate inferences and behaviours that are appropriate for the context in which they occur (Lambon Ralph et al. 2017). In well-practised contexts, in which the relevant information is robustly encoded, conceptual representations need little constraint from semantic control processes to produce the correct response. In contrast, situations requiring the retrieval of weakly-encoded information or uncharacteristic features, and the suppression of strong but currently-irrelevant patterns of retrieval, depend more on control processes to shape semantic retrieval (Jefferies et al. 2020). Converging evidence from neuroimaging, patient and neuromodulation studies suggests that left inferolateral prefrontal cortex, posterior middle temporal gyrus, pre-supplementary motor area and intraparietal sulcus form a semantic control network (SCN); these sites all respond to diverse manipulations of semantic control demands (Jefferies and Lambon Ralph 2006; Hoffman et al. 2010; Jefferies 2013; Lambon Ralph 2014; Nozari and Thompson-Schill 2016; Lambon Ralph et al. 2017; Chiou et al. 2018).

An outstanding question concerns the degree to which the neural mechanisms underpinning semantic control are specialised for this domain. A bilateral “multiple demand” network (MDN), including frontal, parietal, cingulate and opercular brain regions (Duncan and Owen 2000; Duncan 2010; Fedorenko et al. 2013), supports a diverse range of cognitively-demanding tasks, including selective attention, working memory (WM), task switching, response inhibition, conflict monitoring and problem-solving (Fedorenko et al. 2013; Fedorenko 2014; Crittenden et al. 2016; Assem et al. 2020; Diachek et al. 2020). Meta-analyses of neuroimaging studies identify a network for semantic control that partially overlaps with MDN (Figure 3; Noonan et al. 2013; Jackson 2020). However, there also appear to be anatomical differences between these networks: regions supporting semantic control extend into more anterior areas of left inferior frontal gyrus, and posterior middle temporal areas, which are not implicated in executive control more generally. Moreover, SCN shows strong left-lateralisation, in contrast to other aspects of control, which are bilateral or even right-lateralized (Gonzalez Alam et al. 2018; Gonzalez Alam et al. 2019; Jefferies et al. 2020).

Moreover, it is still poorly understood whether semantic control demands are analogous to domain-general control processes. Some studies have argued that there are important differences in the processes supported by MDN and SCN: for example, when semantic category is used as the basis of go-no go decisions, behavioural inhibition is still associated with right-lateralised MD regions, not activation within SCN (Gonzalez-Alam et al., 2018). This suggests that semantic control processes are only recruited when conceptual information itself must be controlled, and not whenever semantic tasks become hard. Semantic control might involve distinct neural processes not shared by the control of action or visual attention, since controlled semantic retrieval draws on heteromodal memory representations and information integration, supported by DMN (Price et al. 2015; Margulies et al. 2016; Price et al. 2016; Pylkkänen 2019; Lanzoni et al. 2020), along with control processes (Davey et al. 2016). The SCN sits at the intersection of DMN and MDN, showing structural and intrinsic functional connectivity to regions in both networks (Davey et al., 2016) and falling between these networks on whole-brain connectivity and functional gradients (Wang et al. 2020): in this way, it might support functional coupling between DMN and MDN in the left-lateralised semantic network. While a few studies have manipulated both linguistic and non-linguistic demands, observing common modulation of the neural response in anterior insula and/or anterior cingulate cortex (Eckert et al. 2009; Erb et al. 2013; Fedorenko et al. 2013; Piai et al. 2013), prior studies failed to match the task structure and task difficulty across these domains. More importantly, we are still lacking knowledge about whether MDN and SCN regions share the same neural coding.

Here, we conducted a pair of fMRI studies to assess the nature of neural signals relating to semantic and domain-general control demands. First, we contrasted parametric manipulations of difficulty for semantic judgements (by varying the strength of association) and verbal working memory (by varying load), to identify sites specifically implicated in semantic and non-semantic control. We matched the task/trial structure and input modality across semantic and non-semantic domains. Next, using pattern classification analyses which examine the multivariate pattern of activation across voxels (Haynes and Rees 2006; Norman et al. 2006; Tong and Pratte 2012; Haynes 2015), we tested which regions in the brain could decode semantic demands and working memory load. Finally, we assessed whether SCN and MD regions could cross-classify difficulty across semantic and non-semantic judgements. In this way, the current study tests the extent to which a shared neural currency underlies both semantic control and working memory load.

## Materials and Methods

### Participants

A group of 32 young healthy participants aged 19∼35 (mean age = 21.97 ±c3.47 years; 19 females) was recruited from the University of York. They were all right-handed, native English speakers, with normal or corrected-to-normal vision and no history of psychiatric or neurological illness. The study was approved by the Research Ethics Committee of the York Neuroimaging Centre. All volunteers provided informed written consent and received monetary compensation or course credit for their participation. The data from one task was excluded for four participants due to head motion, and one additional working memory dataset was excluded due to errors in recording the responses. The final sample included 28 participants for the semantic task and 27 participants for the working memory task, with 26 participants completing both tasks.

### Design

Participants completed two experiments, presented in separate sessions. The first session included four functional scans while participants performed a semantic association task. The second session included three working memory functional scans and a structural scan (see Figure 1 for an example of each task). A slow event-related design was adopted for the two sessions in order to better characterise the activation pattern for each trial. Each trial lasted 9s and each run included 48 trials in the semantic task and 40 trials in the working memory task.

**Figure 1.**
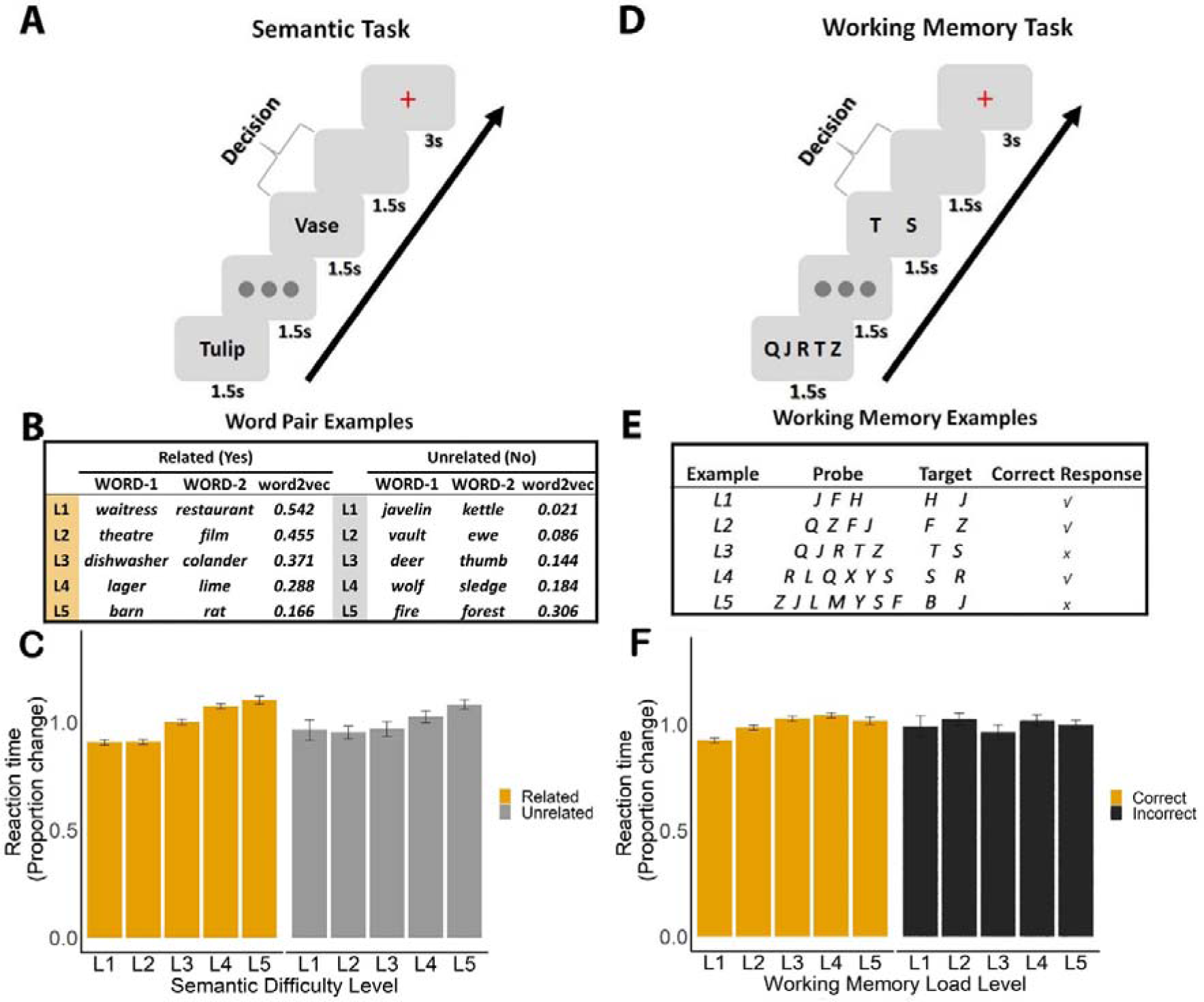
Experiment paradigm and behavioural results. A. Semantic association task; participants were asked to decide if word pairs were semantically related or not. B. Word pair examples for both related and unrelated decisions from one participant, with association strength increasing from Level 1 (L1; little semantic overlap) to Level 5 (L5; high semantic overlap). These trials were assigned to related and unrelated sets of trials on an individual basis for each participant, depending on their decisions, and then split into 5 levels, based on word2vec scores. C. RT for semantic decisions across 5 levels of word2vec for word pairs judged to be related and unrelated. D. Working memory task; participants were asked to decide if two probe letters were presented in a sequence, in any order. E. Working memory load ranged from 3 to 7 items. F. RT for working memory trials across 5 levels of load, for correct and incorrect decisions.

### Semantic association task design

Participants were asked to decide if pairs of words were semantically associated or not. The stimuli were 192 English concrete noun word-pairs. We excluded any abstract nouns and items drawn from the same taxonomic category, so that only thematic links were evaluated in this task (i.e. forest – path or bath – duck; these items are related because they are found or used together). The strength of the thematic link between the items varied parametrically from no clear link to highly related; in this way, participants were free to decide based on their own experience if the words had a discernible semantic link. There were no ‘correct’ and ‘incorrect’ responses: instead, we expected slower response times and less convergence across participants for items judged to be ‘related’ when the associative strength between the items was weak, and for items judged to be ‘unrelated’ when the associative strength between the items was strong (see behavioural below). Overall, there were roughly equal numbers of ‘related’ and ‘unrelated’ responses across participants.

Each trial began with a visually presented word (WORD-1) which lasted 1.5s, followed by a fixation presented at the centre of the screen for 1.5s. Then, the second word (WORD-2) was presented for 1.5s, followed by a blank screen for 1.5s. Participants had 3s from the onset of WORD-2 to judge whether this word pair was semantically associated or not by pressing one of two buttons with their right hand (using their index and middle fingers). During the inter-trial interval (3s), a red fixation cross was presented until the next trial began. Both response time (RT) and response choice were recorded. Participants finished 4 runs of the semantic task, each lasting 7.3 min. Before the scan, they completed a practice session to familiarise themselves with the task and key responses (see Figure 1 for task schematic).

### Semantic stimuli

To quantify the strength of semantic relationships in the association task, distributed representations of word meanings were obtained from the word2vec neural network, trained on the 100 billion-word Google News dataset (Mikolov et al. 2013). In common with other distributional models of word meaning, the word2vec model represents words as high-dimensional vectors with 300 dimensions, where the similarity of two words’ vectors indicates that they appear in similar contexts, and thus are assumed to have related meanings. The word2vec vectors used here were found to outperform other available vector datasets in predicting human semantic judgements in a recent study (Pereira et al. 2016). We defined the strength of the semantic relationship between words using the cosine similarity method. This value was calculated for each word pair presented as a trial, allowing us to characterise the trials on a continuum from strongly related to unrelated.

While word2vec values were higher for trials judged to be semantically related overall (see below), there was considerable variation for both related and unrelated judgements. Since different numbers of items were judged to be thematically related and unrelated across participants, we split related and unrelated trials for each participant into five levels according to their word2vec value, each with the same number of word-pairs. In order to simplify the presentation of the results, the analysis was based on these five levels of word2vec unless otherwise stated. We reasoned that higher word2vec values would be associated with lower task demands for trials judged to be related, and with higher task demands for trials judged to be unrelated.

This was confirmed by behavioural analyses (see below). Word2vec values did not correlate with psycholinguistic variables from N-Watch (Davis 2005), including word length (number of letters: Word1, r = 0.099, p = 0.17; Word2, r = 0.113, p = 0.119), word frequency (Word1, r = 0.033, p = 0.657; Word2, r = 0.111, p = 0.127) or imageability (Word1, r = -0.004, p = 0.958; Word2, r = -0.010, p = 0.901). We also computed a semantic decision consistency index for each word pair by calculating how many participants judged it to be semantically associated (expressed as a proportion of the total participants tested). Word2vec was significantly positively correlated with this consistency value (r = 0.773, p < 0.0001), showing that people were more likely to judge word pairs as related when they had high word2vec values.

### Verbal working memory task

The working memory task had a similar structure to the semantic task (see Figure 1). Each trial began with a letter string (3 to 7 letters) presented at the centre of the screen for 1.5s, followed by a fixation presented for 1.5s. Participants were asked to remember these letters. Next, two letters were shown on the screen for 1.5s.

Participants judged whether both of them had been presented in the letter string by pressing one of two buttons within 3s (participants were told the order of the letters on the screen did not matter). Then a red fixation cross was presented for 3s, until the start of the next trial. Participants completed 3 runs, each containing 40 trials and lasting for 6.1 minutes. The working memory load was manipulated by varying the number of letters memorised in each trial; there were five levels of load from 3 to 7 letters (to match the five levels of word2vec in the semantic task), with 8 trials at each level in each run, presented in a random order. Both response time (RT) and accuracy were recorded, and participants were asked to respond as quickly and accurately as possible.

### Mixed-Effects Modelling of Behavioural Data

Since participants judged different numbers of items to be semantically related and unrelated in the semantic task, mixed-effects modelling was used for the analysis of the behavioural data. This approach is particularly suitable when the number of trials in each condition differs across participants (Mumford and Poldrack 2007; Ward et al. 2013). Semantic association strength (or working memory load) was used as a predictor of the decision participants made (in the semantic task: judgements of whether the words were related or unrelated; in the working memory task: whether the response was correct or incorrect) and, in separate models, how long the reaction time this decision took (i.e., RT). Participants were included as a random effect. The mixed-effects model was implemented with lme4 in R (Bates et al. 2014). We used the likelihood ratio test (i.e., Chi-Square test) to compare models with and without the effect of semantic association strength and working memory load level, in order to determine whether the inclusion of the difficulty manipulations significantly improved the model fit.

### Neuroimaging data acquisition

Imaging data were acquired on a 3.0 T GE HDx Excite Magnetic Resonance Imaging (MRI) scanner using an eight-channel phased array head coil at the York Neuroimaging Centre. A single-shot T2*-weighted gradient-echo, EPI sequence was used for functional imaging acquisition with the following parameters: TR/TE/ = θ 1500 ms/15 ms/90°, FOV = 192 × 192 mm, matrix = 64 × 64, 3 x 3 x 4 mm voxel size, 32 axial slices without a gap. Slices were tilted approximately 30**°** relative to the AC-PC line to improve the signal-to-noise ratio in the anterior temporal lobe and orbitofrontal cortex (Deichmann et al. 2003; Wimmer and Büchel 2019). Anatomical MRI was acquired using a T1-weighted, 3D, gradient-echo pulse-sequence (MPRAGE). The parameters for this sequence were as follows: TR/TE/θ = 7.8s/2.3 ms/20°, FOV = 256 × 256 mm, matrix = 256 × 256, and slice thickness = 1 mm. A total of 176 sagittal slices were acquired to provide high-resolution structural images of the whole brain. The relatively short TE was used to minimise the EPI distortion around ATL. We calculated the temporal signal-to-noise ratio (tSNR) for each participant by dividing the mean of the smoothed time series in each voxel by its standard deviation in each run; we then averaged the tSNR across all runs for the semantic task. These tSNR values were comparable with previous studies (Hoffman et al. 2015; Striem-Amit et al. 2018), and were at acceptable levels (Murphy et al. 2007), although lowest at the anterior temporal pole (mean value: 107.8). Supplementary Figure S8 shows tSNR for a range of ROIs and the full tSNR map in MNI space is available to view online: https://neurovault.org/images/441927/.

### fMRI Data Pre-processing Analysis

Image pre-processing and statistical analysis were performed using FEAT (FMRI Expert Analysis Tool) version 6.00, part of FSL (FMRIB software library, version 5.0.11, www.fmrib.ox.ac.uk/fsl). The first 4 volumes before the task were discarded to allow for T1 equilibrium. The remaining images were then realigned to correct for head movements. Translational movement parameters never exceeded one voxel in any direction for any participant or session. Data were spatially smoothed using a 5 mm FWHM Gaussian kernel. The data were filtered in the temporal domain using a nonlinear high-pass filter with a 100 s cut-off. A two-step registration procedure was used whereby EPI images were first registered to the MPRAGE structural image (Jenkinson and Smith 2001). Registration from MPRAGE structural image to standard space was further refined using FNIRT nonlinear registration (Andersson et al. 2007).

### Univariate Parametric Modulation Analysis

We examined the parametric effect of semantic control demands (i.e. the strength of association between WORD-1 and WORD-2) in the decision phase of the task, using general linear modelling within the FILM module of FSL with pre-whitening turned on. Trials judged to be semantically related (YES trials) and unrelated (NO trials) by participants were separately modelled, using their demeaned word2vec values as the weight, and the RT of each trial as the duration. In addition, we included unmodulated regressors for the trials judged to be related and unrelated, as well as regressors containing WORD-1 and the within-trial fixation between the words. The second fixation interval between the trials was not coded and thus treated as an implicit baseline. Regressors of no interest were included to account for head motion. Three contrasts (related vs. baseline, unrelated vs. baseline, and related vs. unrelated) were defined to examine the effect of semantic control demands on trials judged to be related and unrelated.

The working memory task was analysed in a similar way. Correct and incorrect trials were separately modelled. For correct trials, the parametric effect of difficulty was modelled by including memory load as the weight, and reaction time as the duration of each trial; we also included unmodulated regressors for these trials. In addition, we included three unmodulated regressors: incorrect trials, the first word and the first within-trial fixation. The second fixation interval between the trials was not coded and thus treated as an implicit baseline. Regressors of no interest were included to account for head motion. Two contrasts (correct > baseline and the reverse) were defined to examine how memory load parametrically modulated neural activation in the brain.

For both semantic and WM models, a higher-level analysis was conducted to perform cross-run averaging using a fixed-effects model. These contrasts were then carried forward into the group-level analysis, using FMRIB’s Local Analysis of Mixed Effects 1 + 2 with automatic outlier detection (Beckmann et al. 2003; Woolrich et al. 2004; Woolrich 2008). Unless otherwise noted, group images were thresholded using cluster detection statistics, with a height threshold of z > 3.1 and a cluster probability of p < 0.05, corrected for whole-brain multiple comparisons using Gaussian Random Field Theory. The same threshold was used for both univariate and MVPA analysis. Uncorrected statistical maps are available to view online: (https://neurovault.org/collections/8710/).

### Multivoxel Pattern Analysis

#### Single-trial Response Estimation

We used the least square-single (LSS) approach to estimate the activation pattern for each trial during the decision phase in the two tasks. Each trial’s decision was separately modelled in one regressor and all other trials were modelled together as a second regressor; we also included WORD-1 and the fixation as additional regressors. Pre-whitening was applied. The same pre-processing procedure as in the univariate analysis was used except that no spatial smoothing was applied. This voxel-wise GLM was used to compute the activation associated with each trial in the two tasks. Classification was performed on t statistic maps, derived from beta weights associated with each regressor, to increase reliability by normalizing for noise (Walther et al., 2016).

### Network Selection and Parcellation

We used two complementary multivariate approaches to assess representations of control demands; both network/ROI-based and whole-brain searchlight methods. We used two networks defined from previous studies: the semantic control network (SCN) and multiple-demand network (MDN) (Fedorenko et al. 2013; Jackson 2020). We decomposed these networks into semantic control specific (SCN specific) areas, which did not overlap with MDN; multiple-demand specific (MDN specific) regions, which did not overlap with SCN; and shared control regions identified from the overlap between MDN and SCN. As a comparison, we also examined regions within the semantic network not implicated in control. To identify these regions, we downloaded a semantic meta-analysis from Neurosynth (search term ‘semantic’; 1031 contributing studies; http://www.neurosynth.org/analyses/terms/). Then, we removed regions within this semantic network which overlapped with the two control networks to identify semantic regions predominately associated with semantic representation or more automatic aspects of semantic retrieval, mostly within default-mode network (e.g. in lateral temporal cortex and angular gyrus). All of the voxels within the network maps defined above were included within network-based ROIs.

Intraparietal sulcus was not included because the sequence did not allow us to cover the whole brain for some participants. In total, thirty ROIs were defined; four ROIs in semantic non-control areas, three ROIs in SCN areas, six ROIs in the overlap of MDN and SCN, and seventeen ROIs in MDN specific areas. These thirty ROIs are available online: https://osf.io/bau5c/, see Figure S3A for the four networks.

### Support Vector Regression Analysis

Epsilon-insensitive support vector regression analysis (SVR) (Drucker et al. 1997) was conducted using a linear support vector machine (SVM) (Chang and Lin 2011) and custom code implemented in MATLAB (The MathWorks) (code is available at: https://osf.io/bau5c/). In contrast to conventional support vector machine classification (SVM), the SVR does not depend on categorical classification (i.e., predictions falling on the correct or incorrect side of a hyperplane); instead, it outputs estimations using a regression approach. This approach was used to estimate the difficulty level or cognitive demand for each trial. For each level of difficulty (based on inverse word2vec for semantic trials judged to be related, word2vec for semantic trials judged to be unrelated and memory load in the working memory task), the test and training data were normalized (i.e., mean subtracted and divided by the standard deviation) across voxels within each region of interest (i.e., searchlight, ROI) (Misaki et al. 2010). This allowed an evaluation of the pattern of activity across voxels without contamination from mean signal differences within the searchlight or ROI across the difficulty levels (i.e., the univariate effect) (Misaki et al. 2010; Jimura and Poldrack 2012; Coutanche 2013). The SVR cost parameter was set to 0.001. For each searchlight or ROI, the accuracy of SVR prediction was then calculated within-participant, defined as the z-transformed Pearson’s correlation coefficient between actual and predicted values of the difficulty parameter for the left-run-out data, with the actual difficulty levels ranging from 1 (easy) to 5 (hard) in both tasks. The epsilon parameter in the SVR model was set to epsilon = 0.01 (Jimura and Poldrack 2012).

For each participant, three separate SVR classifiers were trained to decode cognitive demands: these examined the difficulty of semantic trials judged to be related (difficulty maximised for low association trials), the difficulty of semantic trials judged to be unrelated (difficulty maximised for high association trials), and the difficulty of working memory trials (difficulty maximised for highest memory load). We examined generalization of difficulty effects within the semantic domain (i.e. between trials judged to be semantically-related and unrelated). In order to test whether semantic control and executive control share a common neural code, we also performed a series of generalization (cross-task classification) analyses, in which classifiers were trained on each task type (semantic related; semantic unrelated; working memory) and tested on the other task types (i.e. trained on semantic related, tested on working memory), resulting in 4 SVR decoding accuracy types. All classification analyses were performed using a leave-one-run-out cross-validation. SVR decoding was performed using searchlight and ROI approaches.

For searchlight-based analysis, for each voxel, signals were extracted from a cubic region containing 125 surrounding voxels. The searchlight analysis was conducted in standard space. A random-effects model was used for group analysis. Since no first-level variance was available, an ordinary least square (OLS) model was used.

For the network ROI-based analysis, because the number of voxels in the network ROIs varied and differences in ROI size are likely to influence classifier performance, classification analyses were performed by randomly subsampling 200 voxels from each ROI. This process was repeated for 100 iterations for each ROI and subject, with each iteration involving a different random sample of 200 voxels. The 100 iterations in each ROI were averaged into one value, and this value from all ROIs were averaged again for each brain network.

## Results

### Behavioural Results

Overall, equal numbers of word pairs were judged to be related or unrelated by the participants (mean ratio: 0.491 vs. 0.495, χ2(1) = 0.00021, p > 0.995). Linear mixed effects models examined whether associative strength and working memory load were reliable predictors of behaviour. We found that both the strength of the semantic association (word2vec value) and working memory load successfully manipulated task difficulty. For the semantic task, the continuous word2vec value was positively associated with a higher probability that participants would identify a semantic relationship between the words ( 2(1) = 2421.3, p < 0.001) using a logistic χ regression approach. When word pairs were grouped into 5 levels according to their word2vec value, the relationship was still significant ( 2(1) = 2467.8, p < 0.001).

Since we used a continuous manipulation of associative strength, and there is no categorical boundary of word2vec values which can capture the trials reliably judged to be related and unrelated, we were not able to compute a traditional error score for the semantic task. We expected that for those word-pairs judged to be related in meaning, higher word2vec values would facilitate semantic decision-making. For these trials, the pattern of semantic retrieval required by the task (i.e. the identification of a linking context) is likely to be well-supported by dominant information in long-term memory. Since the linking context is highly accessible on these trials, there is less uncertainty about the relevant response, and potential conflict between the response options is reduced. In contrast, when items are judged to be semantically related even when they have less semantic overlap as assessed by word2vec, it is thought that control processes must be engaged to shape activation within the semantic store; this is because a dominant linking context is not readily available in long-term memory. In this situation, task-irrelevant but more dominant semantic associations to the two words may need to be suppressed and there is likely to be more uncertainty about the decision. For trials in which words are judged to be unrelated in meaning, the effect of strength of association is expected to have the opposite effect on task difficulty. When the two items have very different meanings and are not remotely connected to each other, it is relatively easy to decide that they are not semantically associated; low word2vec values should be associated with lower decisional uncertainty. In contrast, when participants decide that two words are unrelated even when they are somewhat linked according to word2vec, the semantic decision is expected to be more difficult, with greater uncertainty or response conflict emerging from their partial relationship. Participants may need to recruit control processes to overcome this conflict or uncertainty.

Mean RT for each level is presented in Figure 1C, separately for related (YES) and unrelated (NO) decisions. To examine how association strength level modulated RT for trials judged to be related and unrelated, we performed linear mixed effects analyses with participant as a between-subject variable and association level as a within-subject variable. This revealed a significant effect of level of association strength for both related and unrelated decisions. Association strength level was negatively associated with RT (χ2(1) = 146.6, p < 0.001) for related trials and positively associated with reaction time for unrelated trials (χ = 58.668, p < 0.001). It was more difficult for participants to retrieve a semantic connection between two words when strength of association was lower; on the contrary, it was easier for them to decide there was no semantic connection between word pairs with low word2vec values.

For the working memory task, the proportion of correct responses was 84.8%, when all memory load levels were considered. The more items to be maintained or manipulated in working memory, the more difficult the trial was expected to become. A logistic regression showed that higher working memory load was associated with lower accuracy (χ2(1) = 112.4, p < 0.001). A further linear mixed effects model with participant as a between-subject variable and memory load as a within-subject variable revealed a significant positive relationship between load level and RT for correct responses (χ2(1) = 39.826, p < 0.001).

Lastly, a two-way repeated-measures ANOVA was conducted examining the effects of task condition (semantic related, unrelated and working memory correct) and difficulty level (five levels per task) on the proportional change in RT for each difficulty level of the task, relative to the average RT for each condition. The results showed a significant interaction between conditions and difficulty levels (F(5.395,134.881) = 8.329, p < 0.001, Greenhouse-Geisser corrected), along with a main effect of difficulty level (F(3.134, 78.346) = 53.262, p < 0.001, Greenhouse-Geisser corrected). Together, these results suggest that association strength and memory load successfully manipulated task difficulty, with the semantic task showing a stronger influence on RT than WM load.

### fMRI Results

#### The Parametric Effect of Word2vec on Brain Activation

We identified brain areas showing an increase or decrease in activation as a function of association strength (using the continuous word2vec scores). It was harder for participants to decide that items were semantically related when they were weakly associated; consequently, we would expect stronger responses in semantic control and multiple demand regions for these trials. It was also harder for participants to decide that items were semantically unrelated when they had greater word2vec values; therefore we would expect opposite effects of word2vec for related and unrelated trials in brain regions supporting demanding semantic decisions. The direct comparison of word2vec effects for semantically-related and unrelated decisions can identify brain areas responding to semantic similarity but not difficulty, while the combination of negative effects of word2vec for related decisions and positive effects of word2vec for unrelated decisions can identify brain regions that respond to the difficulty of semantic decisions, without a confound of semantic relatedness.

For related trials, weaker associations elicited greater activity in regions linked to semantic control in previous studies, including left inferior frontal gyrus (IFG), left middle frontal gyrus (MFG), superior frontal gyrus (SFG) and left posterior middle temporal gyrus (pMTG); see Figure 2A. Similarly, when participants decided that items were unrelated, there was stronger activation in left inferior frontal gyrus, middle frontal gyrus, superior frontal gyrus and frontal orbital cortex (FOC) when these items had higher word2vec scores; see Figure 2B.

**Figure 2.**
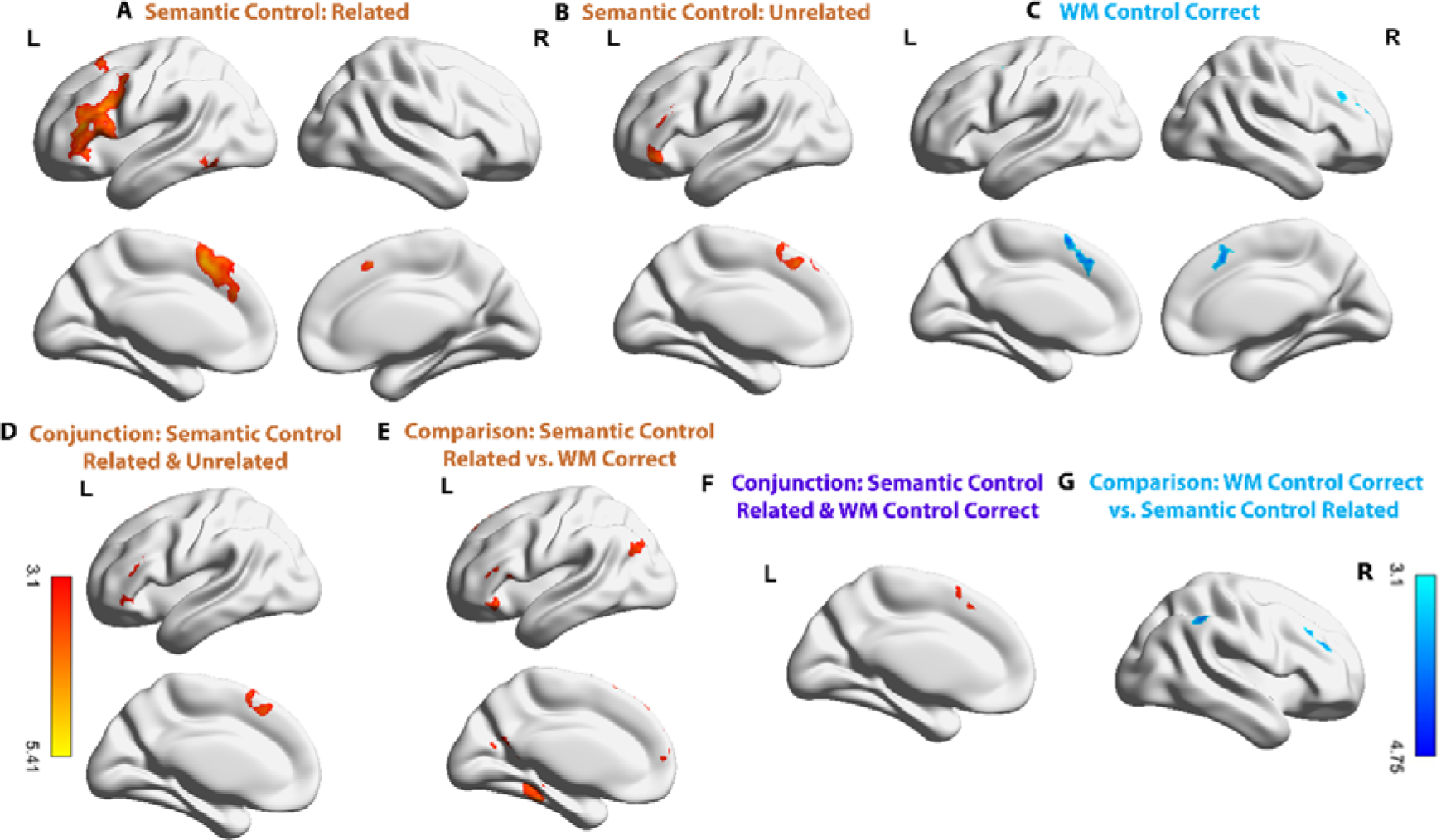
Univariate results with cluster thresholded at Z = 3.1, p = 0.05. A. Parametric modulation effect of associative strength for trials judged to be semantically related. B. Parametric modulation effect of associative strength for trials judged to be unrelated. C. Parametric modulation effect of working memory for correct trials. D. The conjunction of semantic control parametric effects across trials judged to be related and unrelated (i.e. negative word2vec for related trials and positive word2vec for unrelated trials). E. Areas showing a stronger parametric effect of control demands for semantic judgements (negative effect of word2vec for related trials) compared to working memory (effect of memory load for correct trials). F. The conjunction of the parametric modulation effect for semantic control (from related trials) and working memory load (correct trials). G. A larger parametric modulation effect for working memory load (correct trials) compared with semantic control demands (negative effect of word2vec on semantically-related trials). There are no additional clusters within brain views not shown for each contrast.

We investigated common and distinct effects of semantic control demands across trials classified as related and unrelated. A conjunction analysis revealed that rejecting strongly associated word pairs and accepting weakly associated word pairs recruited common semantic control regions including left inferior frontal gyrus, middle frontal gyrus, superior frontal gyrus and frontal orbital cortex, see Figure 2D. There were no significant differences in the parametric effects of semantic control demands or semantic relatedness for trials judged to be related and unrelated in a direct contrast. There were also no common effects of semantic similarity (i.e. positive effects of word2vec that were shared across related and unrelated decisions).

In addition, although we could not compute task accuracy in our main analysis (since we manipulated strength of association in a continuous way, and participants were asked to split this distribution into related and unrelated trials), a supplementary control analysis removed trials with unexpected word2vec scores, given the decision that was made. An additional regressor was included to capture trials judged to be related even though they had particularly low word2vec values (bottom 25% of word2vec values), and trials judged to be unrelated that had particularly high word2vec values (top 25% of word2vec values). The results were very similar to the analysis above; see Supplementary Figure S1A.

#### The Parametric Effect of Working Memory Load on Brain Activation and the Comparison with Semantic Control

For correct working memory trials, a significant parametric effect of memory load was found in right middle frontal gyrus, frontal pole (FP) and superior frontal gyrus consistent with previous studies in which higher working memory loads elicited greater activity in distributed bilateral areas within the multiple-demand network (MDN); see Figure 2C.^1^

We performed further analyses to establish the common and distinct parametric effects of semantic control demands and working memory load. We compared correct working memory trials and word pairs judged to be semantically-related, since participants made YES decisions in both situations. Since semantic relatedness was varied in a continuous fashion while working memory load was manipulated across five levels, we first divided the semantically related trials into five difficulty levels according to their word2vec values, with lower word2vec corresponding to harder trials (re-analysis of the univariate activation for the semantic task using these five levels replicated the findings above and obtained highly similar results, see Supplementary Figure S1B). To simplify the following univariate and multivariate results focussed on the comparison of the semantic and working memory tasks, we used five levels of difficulty or association strength for the thematically related and unrelated decisions, unless otherwise mentioned.

A conjunction analysis showed a significant overlap between semantic control demands and working memory load in superior frontal gyrus and pre-supplementary motor area (pre-SMA); see Figure 2F. Direct contrasts of these semantic and non-semantic difficulty effects revealed stronger effects of difficulty in the working memory than the semantic task in right-lateralized regions mainly within the multiple-demand network, including right middle frontal gyrus, frontal pole and supramarginal gyrus (SMG); Figure 2G. There was a greater effect of semantic control demands in distributed areas in the left hemisphere, including IFG, frontal orbital cortex, superior frontal gyrus, lateral occipital cortex (LOC), precuneus, hippocampus, parahippocampal gyrus and temporal fusiform, consistent with previous observations that semantic control is strongly left-lateralized; Figure 2E. A supplementary ROI-based analysis using percent signal change to directly compare the parametric effect of difficulty against implicit baseline in the two tasks showed that the task differences in most of the clusters in Figure 2E and Figure 2G were driven by increased responses to more difficult trials, and not solely by negative parametric effects of difficulty in the other task (see detailed information in Supplementary Figure S2B).

#### Identifying the Neural Coding of Semantic and Working Memory Demand Using a Searchlight Approach

In order to test whether the same neural code supported semantic control demands and working memory load, we examined classification of control demands (task difficulty) in each task, and cross-classification of difficulty across tasks using a whole-brain searchlight approach. For the semantic task, word2vec (as a measure of relatedness) and difficulty (as assessed by behavioural performance) show an opposite relationship for trials judged to be related and unrelated. We therefore reasoned that a classifier sensitive to control demands would show a positive correlation between actual and predicted control demands when trained on related trials (which had low word2vec values for more difficult trials) and then tested on unrelated trials (which had high word2vec values for more difficult trials), or vice versa. In contrast, brain regions showing a negative correlation across these trial types would be sensitive to the associative strength of the presented items, irrespective of the subsequent judgement. Moreover, brain regions able to cross-classify difficulty between semantic and working memory tasks are sensitive to domain-general control demands.

After controlling for univariate activation (see Methods), we found difficulty could be decoded for semantically-related trials in lateral and medial frontal and parietal areas, bilaterally, as well as left posterior middle temporal gyrus, see Figure 3A. We also found significant decoding of difficulty for semantically-unrelated trials in similar areas, see Figure 3B. Finally, we searched for brain regions that supported cross-classification of difficulty across semantically-related and unrelated trials (training on one condition and testing on the other). Significant positive cross-classification was identified in distributed regions including left inferior and middle frontal gyrus,bilateral superior frontal gyrus/paracingulate gyrus, left posterior middle temporal lobe, bilateral lateral occipital cortex/angular gyrus, see Figure 3D. These sites were sensitive to semantic difficulty irrespective of strength of association, while there were no significant clusters showing negative correlation in the cross-decoding between related and unrelated trials, suggesting our classifiers were not sensitive to semantic relatedness.

**Figure 3.**
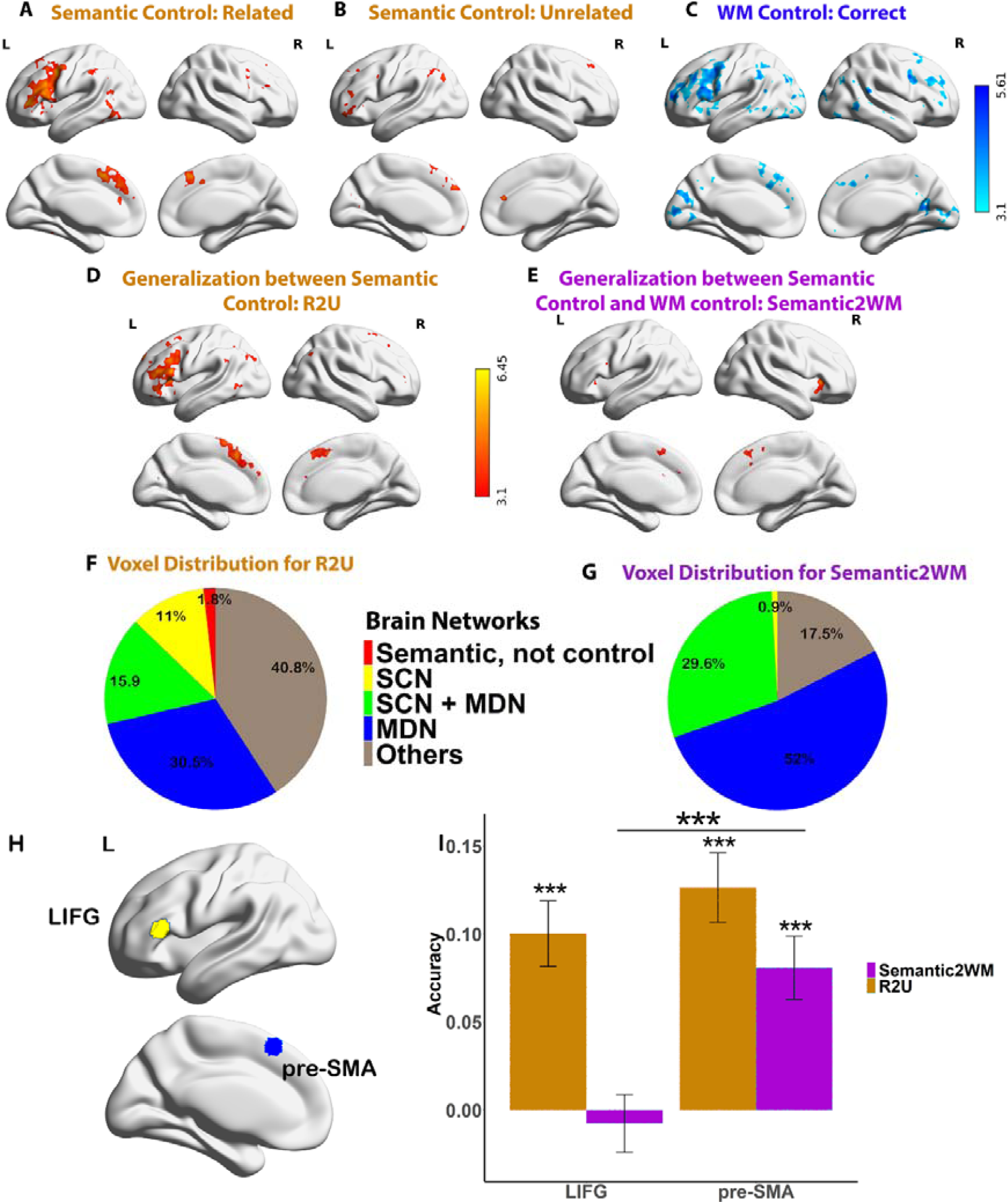
SVR decoding of control demands. A. Brain regions representing control demands for semantic trials judged to be related. B. Brain regions representing control demands for semantic trials judged to be unrelated. C. Brain regions representing working memory load. D. Brain regions with significant cross-classification of difficulty between semantic trials judged to be related and unrelated. E. Brain regions with significant cross-task classification of difficulty. F. Voxel distribution in Figure 3D across regions identified as (i) semantic not control, (ii) within the semantic control network (SCN) but outside multiple-demand cortex, (iii) within both semantic control and multiple-demand networks (SCN+MDN), and (iv) falling in multiple-demand regions not implicated in semantic cognition (MDN). Voxels showing significant cross-classification outside these networks are also shown (Others); 9716 voxels in total. G. Voxel distribution in Figure 3E across semantic not control, SCN only, SCN+MDN, MDN only and other networks, 933 voxels in total. H. ROIs definition example. The SCN ROI was defined for each participant individually using their peak coordinates in the univariate contrast between parametric effects of control demands in semantic judgements compared to working memory load in the left inferior frontal gyrus (LIFG, a cube with 125 voxels, mean MNI coordinates across the sample: X = -50, Y = 28, Z = 12). The MDN ROI was defined using each participant’s peak coordinates in the univariate conjunction analysis of the parametric effects of semantic control demands (for related trials) and working memory load (correct trials) in the presupplementary motor cortex (pre-SMA, a cube with 125 voxels, mean MNI coordinates across the sample: X = -6, Y = 22, Z = 52). I. There was only significant cross-task decoding of difficulty across semantic and WM tasks in the pre-SMA, but no such effect in the LIFG, and the cross-task decoding accuracy was significant higher in pre-SMA than in LIFG. Both ROIs supported cross-trial decoding of difficulty between related and unrelated trials within the semantic domain, and there was no significant difference in decoding accuracy between pre-SMA and LIFG. All p values were Bonferroni corrected. ***p < 0.001/2. **p < 0.01/2. *p < 0.05/2.

Brain regions that coded for working memory load were found in frontal, parietal, temporal as well as visual cortex, bilaterally (see Figure 3C). Compared to the neural underpinnings of semantic control, which were strongly left-lateralised, the multivariate effect of working memory load was bilateral. There was significant cross-task classification between semantic and working memory tasks in bilateral insula, pre-supplementary motor area and left precentral gyrus, see Figure 3E. Most of these voxels fell within the SCN+MDN (29.6%) and MDN (52%), and few were within SCN (0.9%), see Figure 3G. This result suggests that SCN regions do not support a shared neural coding between semantic and non-semantic control demands, while MDN regions (including those overlapping with SCN) show common patterns across manipulations of semantic and working memory control demands. In contrast, cross-classification of difficulty between related and unrelated semantic decisions overlapped with SCN-only regions (9716 voxels in total, see Figure 3F).

Finally, to establish whether regions in MDN showed higher cross-task decoding accuracy than SCN regions, we conducted an ROI analysis. This analysis also established whether SCN regions can decode non-semantic as well as semantic task demands when a less stringent threshold is applied (i.e. when no multiple comparison correction for whole-brain analysis is applied). ROIs (cubes containing 125 voxels) were defined from the univariate parametric analysis and from published network maps (see Supplementary Materials Figure S3). We defined the SCN ROI for each participant using their peak response to the contrast of parametric effects of control demands for semantic judgements compared to working memory load in the left inferior frontal gyrus (LIFG). We defined the MDN ROI using each participant’s peak coordinate for the conjunction of parametric effects for semantic control (from related trials) and working memory load in the presupplementary motor cortex (pre-SMA). We extracted the decoding accuracy for each participant in these two ROIs and compared them using ANOVA. There was a significant interaction between network ROI and cross-classification type (cross-classification of difficulty across related and unrelated trials within the semantic domain vs. cross classification of semantic control demands and working memory load; F(1,25) = 5.38, p = 0.029).

There were also significant main effects of cross-classification type (F(1,25) = 21.98, p < 0.001) and ROI (F(1,25) = 14.66, p = 0.001). Simple t-tests revealed that there was significant cross-task decoding in pre-SMA (p < 0.001), but no such effect in LIFG (p = 1). As expected, cross-task decoding accuracy was significantly higher in pre-SMA than in LIFG (p < 0.001). Both ROIs supported cross-condition decoding of difficulty within the semantic domain (i.e. between trials judged to be related and unrelated; p < 0.001), and there was no significant difference in decoding accuracy between pre-SMA and LIFG (p = 0.459). All p values are adjusted by Bonferroni correction.

These results demonstrate the critical role of SCN regions in coding for semantic control demands, alongside the role of MDN (in MDN-only and MDN+SCN regions) in representing the difficulty of both semantic and non-semantic control demands in a common neural code.

To further check the robustness of our conclusions, we examined classification of control demands in each task, and cross-classification of difficulty across tasks, within pre-defined MDN and SCN networks. We performed a series of SVR decoding analyses in ROIs selected to fall within the following areas: (i) sites within the semantic network but not implicated in control; (ii) SCN (defined as voxels within the semantic control network identified by Noonan et al. (2013) and updated by Jackson (2020), and yet outside the MDN); (iii) regions common to both SCN and MDN; (iv) MDN (defined as voxels within the multiple-demand network identified by Fedorenko et al. (2013), and not within the SCN. In a control analysis, we also randomly subsampled 200 voxels in each ROI in each network. Details are provided in Supplementary Figures S3 to S4. These analyses support our key conclusions: cross-classification of difficulty between semantic and WM tasks was only found in MDN-only and SCN+MDN regions, not in SCN-only or semantic not control network regions; and there was a decreasing pattern in cross-task decoding and the representation of working memory load from shared MDN+SCN regions, through SCN to semantic regions not implicated in control. However, the difficulty of both tasks could be individually decoded in all four networks (see below for discussion).

## Discussion

This study parametrically manipulated the difficulty of semantic and verbal working memory judgements to delineate common and distinct neural mechanisms supporting control processes in these two domains. Across two experiments, we investigated the brain’s univariate and multivariate responses to different manipulations of difficulty: in a semantic relatedness task, we varied the strength of association between probe and target words, while in a verbal working memory task, we manipulated the number of items to be maintained (working memory load).

Retrieving semantic links between weakly associated words is known to elicit stronger activation within the “semantic control network” (SCN) (Noonan et al. 2013; Jackson 2020), while higher loads in working memory are associated with greater responses within the “multiple demand network” (MDN) (Fedorenko et al. 2013) – particularly, left-lateralised parts of this network for verbal working memory (Emch et al. 2019). This comparison is therefore ideal to establish similarities and differences in the neural basis of these forms of control, with any divergence unlikely to be accounted for by the use of language (as both tasks were verbal in nature). We obtained convergent evidence across analyses for both common and distinct neural responses to difficulty across networks. Dorsolateral prefrontal cortex and pre-supplementary motor area (within MDN) showed a common response to difficulty across tasks; in decoding analyses, MDN showed common patterns of activation across manipulations of semantic and non-semantic demands, and cross-classification of difficulty across tasks. In contrast, left inferior frontal gyrus within SCN showed an effect of difficulty that was greater for the semantic task; moreover, there was no shared neural coding of cognitive demands in SCN regions, consistent with the view that semantic control has a neural basis distinct from other cognitive demands beyond the semantic domain.

The semantic control network, encompassing left inferior frontal gyrus and posterior middle temporal gyrus, is known to activate across a wide range of manipulations of semantic control demands – including a stronger response for weak associations, ambiguous words, and multiple distractors (Noonan et al. 2013; Davey et al. 2016; Jackson 2020). Since these regions are implicated in semantic cognition, as well as in control processes, one point of contention is the extent to which semantic retrieval per se, which is potentially increased in more demanding conditions, can explain this pattern of results. A unique strength of this study is that we can distinguish the impacts of semantic control and within-trial semantic similarity through the comparison of difficulty in trials judged by participants to be related and unrelated.

This is because semantic similarity has opposite effects of difficulty in these two sets of trials: when participants decide there is a semantic link between two words, more control is needed to make this link when the words are more weakly associated; in contrast, when participants decide there is no semantic link between two words, more control is needed for this decision when the words are more strongly associated. The univariate analyses found equivalent effects of difficulty in left inferior frontal gyrus for trials judged to be related and unrelated; consequently, we can conclude this site is sensitive to the difficulty of semantic decisions and not strength of association per se. This is exactly the pattern that we would expect for brain regions implicated in semantic control but not long-term conceptual similarity.

In addition to investigating the involvement of SCN and MDN in semantic and non-semantic tasks differing in difficulty, we examined the characteristics of cognitive control in multivariate analyses of activation patterns using a whole-brain searchlight approach and SVR decoding for the first time. Our results revealed significant information about semantic demands within both SCN and MDN; however, cross-classification of control demands across related and unrelated semantic trials identified regions within SCN that lie beyond MDN, while cross-classification of control demands across semantic and WM tasks identified MDN regions – both regions that overlap with SCN, and other MDN regions that lie beyond the semantic network. These findings point to functional heterogeneity across control network regions (Dixon et al. 2018). Though previous studies revealed that distributed areas in the left lateral frontal, medial frontal, lateral temporal and parietal regions support the representation of semantic relatedness (Mahon and Caramazza 2010), our decoding generalization analysis provided strong evidence that semantically-related and unrelated judgements share the same neural code relating to difficulty in SCN (despite opposing effects of semantic similarity). We additionally demonstrated that working memory load is reflected in the activation patterns of SCN as well as MDN; both networks are sensitive to the difficulty of both semantic and non-semantic tasks: this places limits on observations that language and domain-general control demands are reliant on distinct neural substrates (Fedorenko 2014; Blank and Fedorenko 2017; Diachek et al. 2020; Fedorenko and Blank 2020). Importantly, we were not able to decode difficulty across tasks in SCN (when training on working memory load and testing on semantic association strength or vice versa), even in an ROI analysis, suggesting that the multivariate neural codes relating to the difficulty of semantic and working memory judgements may be distinct in SCN.

In supplementary network-based decoding analyses, we also examined the decoding of task demands within parts of the semantic network not implicated in control processes, primarily regions within default mode network (DMN; including anterior lateral temporal cortex and angular gyrus). Decoding of task difficulty was less accurate in DMN than in control networks – yet semantic regions not implicated in control were still able to decode difficulty across tasks. The DMN has long been considered a ‘task-negative’ network, only engaged when the brain is not occupied by an externally-presented task, and associated with internally-oriented cognitive processes such as mind-wandering, memory retrieval and future planning (Buckner and DiNicola 2019). DMN regions typically show deactivation relative to rest during challenging tasks (Raichle et al. 2001; Raichle 2015). However, our multivariate analysis examined normalized activation across voxels within each region of interest (i.e., searchlight, ROI) (Misaki et al. 2010; Jimura and Poldrack 2012; Coutanche 2013); consequently, greater mean deactivation in hard versus easy tasks is unlikely to be the basis for the decoding results in DMN. Instead, task difficulty may have related to specific patterns of activation and/or deactivation in this network, for example, reflecting the way that DMN changes its pattern of connectivity to suit the ongoing task demands (Cole et al. 2013). Research indicates that semantically-relevant regions of DMN show increased connectivity to executive cortex during control-demanding semantic tasks (Krieger-Redwood et al. 2015). Moreover, beyond the semantic domain, DMN shows dynamically changing patterns of connectivity with other brain networks, including those implicated in control and attention, as working memory load is varied (Vatansever et al. 2015). In line with the view that DMN may play a more active role in even demanding aspects of cognition, recent studies show that multivariate patterns in both DMN and MDN track goal information instead of conceptual similarity (Wang et al. 2020), and that DMN represents broad task context (Wen et al. 2020). The current results are therefore consistent with growing evidence that DMN can contribute to controlled as well as more automatic aspects of cognition, even as it deactivates (Elton and Gao 2015; Raichle 2015; Vatansever et al. 2015; Vatansever et al. 2017).

As proposed by Duncan (2010, 2013, 2016), MDN captures an abstract code relating to the difficulty of decisions across multiple domains. Frontal-parietal regions in MDN have been shown to flexibly represent goal-directed information, including visual, auditory, motor, and rule information, to support context-appropriate behaviour (Cole et al. 2013; Crittenden et al. 2016; Woolgar et al. 2016; Bhandari et al. 2018).

However, the current study, to our knowledge, is the first to test whether semantic and non-semantic verbal demands share a common neural currency in the brain via cross-classification analyses. Converging evidence from both ROI/Network and searchlight-based analyses revealed that only regions in MDN (including the overlap with SCN) could cross-classify task difficulty across domains. This observation is noteworthy given previous proposals that the “language” network is largely distinct from MDN regions in LIFG (Fedorenko and Blank 2020); we find that additional regions are recruited to support semantic control, in line with this view, but that MDN regions are also recruited in these circumstances, giving rise to functional overlap.

Our results suggest that SCN diverges from this pattern in important ways. There was no such common currency in SCN-specific regions or non-control semantic areas. Semantic cognition is thought to emerge from heteromodal brain regions (Lambon Ralph et al. 2017); in line with this, a heteromodal control network which only partially overlaps with MDN has been shown to support the retrieval of both verbal and non-verbal information (Krieger-Redwood et al. 2015). Semantic control processes could regulate the activation of semantic features, thought to draw on unimodal systems supporting vision, audition and action etc. – for example, when linking dog to beach, activation might be focussed on running, swimming and digging actions, as opposed to the physical features of a dog (such as its ears and tail).

These features are thought to rely on interactions between the heteromodal ‘hub’ within anterolateral temporal cortex and ‘spoke’ systems in unimodal cortex; consequently, semantic control processes could bias activation towards relevant spoke systems, resulting in a more relevant response within the heteromodal hub when these features are distilled into a coherent meaning (Jackson et al. 2019; Zhang et al. 2020). This process could be largely analogous to the way that MDN regions are thought to bias processing towards task-relevant inputs or sensory features. However, our observation that there are semantic-specific control processes beyond MDN is consistent with the view that this mechanism is supplemented by separate semantic control representations that interface with the long-term conceptual store. This evidence allows us to reject the account that semantic control demands are exactly analogous to other types of cognitive demand.

Contemporary accounts of brain organization suggest that neural function is organized along a connectivity gradient from unimodal regions of sensorimotor cortex, through executive regions to transmodal default mode network (Margulies et al. 2016; Huntenburg et al. 2018). Wang et al. (2020) suggested this gradient can capture the orderly transitions between MDN, SCN and DMN in semantic processing. Given that SCN has greater proximity to DMN than MDN along this principal gradient of connectivity, this network might be able to more efficiently select, retrieve and act on semantic information stored in heteromodal DMN regions. Our results showed a decreasing pattern in cross-task decoding and the representation of working memory load from MDN and shared MDN+SCN regions, through SCN to semantic regions not implicated in control (see Supplementary Materials Figure S3) – with this series of networks following the principal gradient (Wang et al. 2020). In a similar way, González-García et al. (2018) found regions in DMN and MDN have similar representational formats relating to prior experience, and occupy adjacent positions on the principal gradient.

One limitation of the current study was that different metrics (strength of association and working memory load) were used to manipulate difficulty across the semantic and working memory tasks, and it is difficult to directly compare these manipulations. The WM task was associated with faster responses, perhaps because word reading takes longer than letter identification, but RT reading times are not necessarily relevant to the activation of control networks. Similarly, the effect of strength of association had a larger effect on RT than working memory load, although RT does not provide a direct measure of cognitive control demands. Our task design focussed on manipulating task demands in the verbal domain when semantic cognition was or was not required, and the tasks were similar in their visual presentation and in the button-press response. Future studies could manipulate semantic and non-semantic tasks in more directly comparable ways, for example by presenting strong vs. weak distractors or more vs. less information to support a specific decision. A better match in difficulty across tasks might result in further cross-task classification results, extending beyond the regions identified here.

In summary, univariate and multivariate pattern analyses provide strong evidence that semantic control demands and working memory load recruit both common and distinct processes in the multiple-demand and semantic control networks, respectively. Though semantic demand and domain general demand are represented in both control networks, there was only shared neural coding of difficulty across tasks in MDN, and different neural coding of control demands in SCN. These findings indicate SCN and MDN can be dissociated according to the information that they maintain about cognitive demands.

## Acknowledgements

This work was sponsored by the European Research Council (Project ID: 771863 - FLEXSEM Project).

## Author contributions

Z.G. and E.J. designed the experiment. Z.G. performed the study. Z.G. analysed the data.

Z.G., E.J. wrote the original manuscript, L.Z. R.C. J.S. M.A.L.R. reviewed and edited the manuscript.

## Footnotes

1 A supplementary analysis, thresholded at Z > 2.6, revealed a more distributed neural substrate for working memory load including bilateral middle frontal gyrus, precentral gyrus and occipital fusiform cortex; see Supplementary Figure S2A.

