## Supplementary Materials for "Distinct and Common Neural Coding of Semantic and Non-semantic Control Demands"

**The Parametric Effect of Word2vec on Brain Activation for ‘Correct’ Related and Unrelated Trials**

In this analysis, we removed trials that had relatively unexpected word2vec scores, given the decision that was made. These trials were expected to include the majority of any responses made in error. An additional regressor was included to capture trials judged to be related even though they had particularly low word2vec values (bottom 25% of word2vec values), and trials judged to be unrelated that had particularly high word2vec values (top 25% of word2vec values).

The parametric modulatory effect for ‘correct’ related and unrelated conditions resembled the results we reported in the main manuscript. For related trials, weaker associations elicited greater activity in regions linked to the semantic control network in the left hemisphere, including left inferior frontal gyrus (IFG), left middle frontal gyrus (MFG), superior frontal gyrus (SFG) and left posterior middle temporal gyrus (pMTG); see Figure S1A. Similarly, when participants decided that items were unrelated, there was stronger activation in left superior frontal gyrus and left frontal orbital cortex (FOC) for word-pairs with higher word2vec scores, see Figure S1A. The direct comparison between related and unrelated parametric modulation effects did not identify any clusters, which is also consistent with the results in the main manuscript.

**The Parametric Effect of Word2vec on Brain Activation Using Five levels of Difficulty**

Since semantic relatedness was varied in a continuous fashion while working memory load was manipulated across five levels, an additional analysis was performed to exclude the possibility that task differences in the effects of difficulty were driven by the larger range of semantic relatedness values. To equate the tasks, we first divided the semantically related trials into five difficulty levels according to their word2vec values, with lower word2vec corresponding to harder trials and then re-analysed the univariate parametric effect of semantic difficulty using these five levels. This replicated the findings in the main text; see Figure S1B.

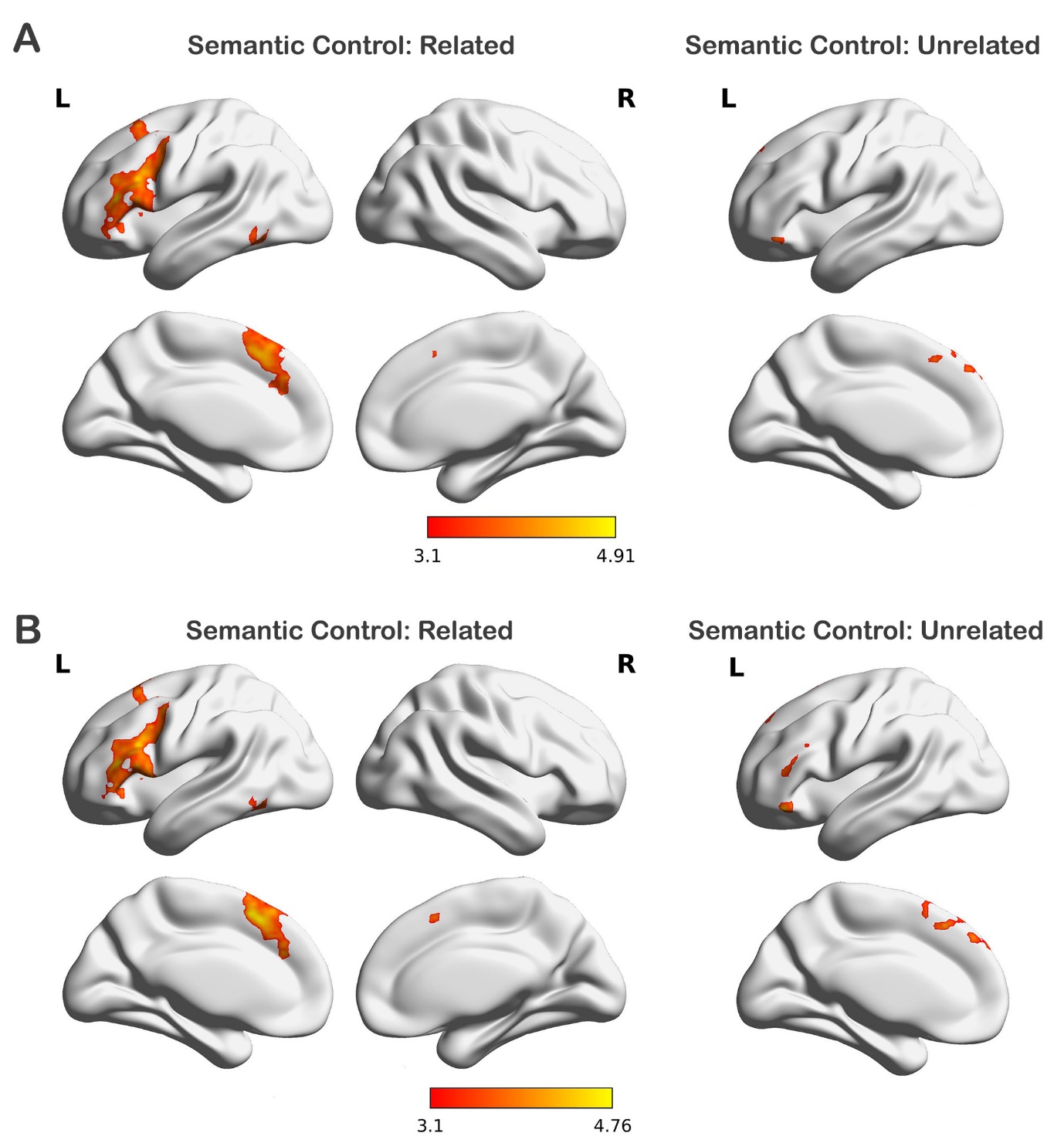

**Figure S1.** Univariate parametric effects analyses while separately modelling ‘correct’ and ‘error’ trials. A. Parametric modulation effect of semantic association strength for the correct related trials (left) and correct unrelated trials (right). B. Parametric modulation effect of semantic difficulty for the related and unrelated trials using five discrete difficulty levels (from one to five), respectively.

**The Parametric Effect of Working Memory Load on Brain Activation and the Comparison with Semantic Control**

In the main text, the group analyses were conducted using cluster detection statistics with a threshold Z > 3.1. In order to avoid type II errors, we also report the results using a lower threshold of Z > 2.6. A significant parametric modulation effect of working memory load was found in more distributed areas within the multiple-demand network (MDN), including right middle frontal gyrus (MFG), frontal pole (FP), and superior frontal gyrus (SFG), and also left middle frontal gyrus (MFG) and precentral gyrus, see Figure S2A.

To establish if the significant task differences shown in the main text (the semantic over working memory difficulty effects in Figure 2E, and the reverse effects in Figure 2G) were driven by positive or negative parametric modulation effects, we extracted the percentage signal change values in each cluster for each task, and conducted a two-way repeated-measures ANOVA including task (semantic related and working memory correct trials) as well as ROI (left inferior frontal gyrus (IFG), left frontal orbital cortex (FOC), medial prefrontal cortex (mPFC), precuneus and temporal fusiform (TempFus) from Figure 2E; and right middle frontal cortex (MPFC) and lateral occipital cortex (LOC) from Figure 2G). Mauchly’s Test of Sphericity indicated that the assumption of sphericity had been violated (χ^2^ = 74.463, p < 0.0001), and therefore, a Greenhouse-Geisser correction was used. There was a significant interaction between ROI and task (F(3.559, 88.984) = 20.128, p < 0.0001). Post-hoc t-tests found a significant difference in each ROI (p < 0.001), see Figure S2B. Further t-tests against the implicit baseline revealed stronger activation for more difficult semantic trials in left inferior frontal gyrus (p < 0.001), frontal orbital cortex (p < 0.001), LOC (p = 0.028), and temporal fusiform (P = 0.03). Right middle frontal gyrus showed stronger activation with higher working memory load (p < 0.001). All p values were adjusted by Bonferroni correction.

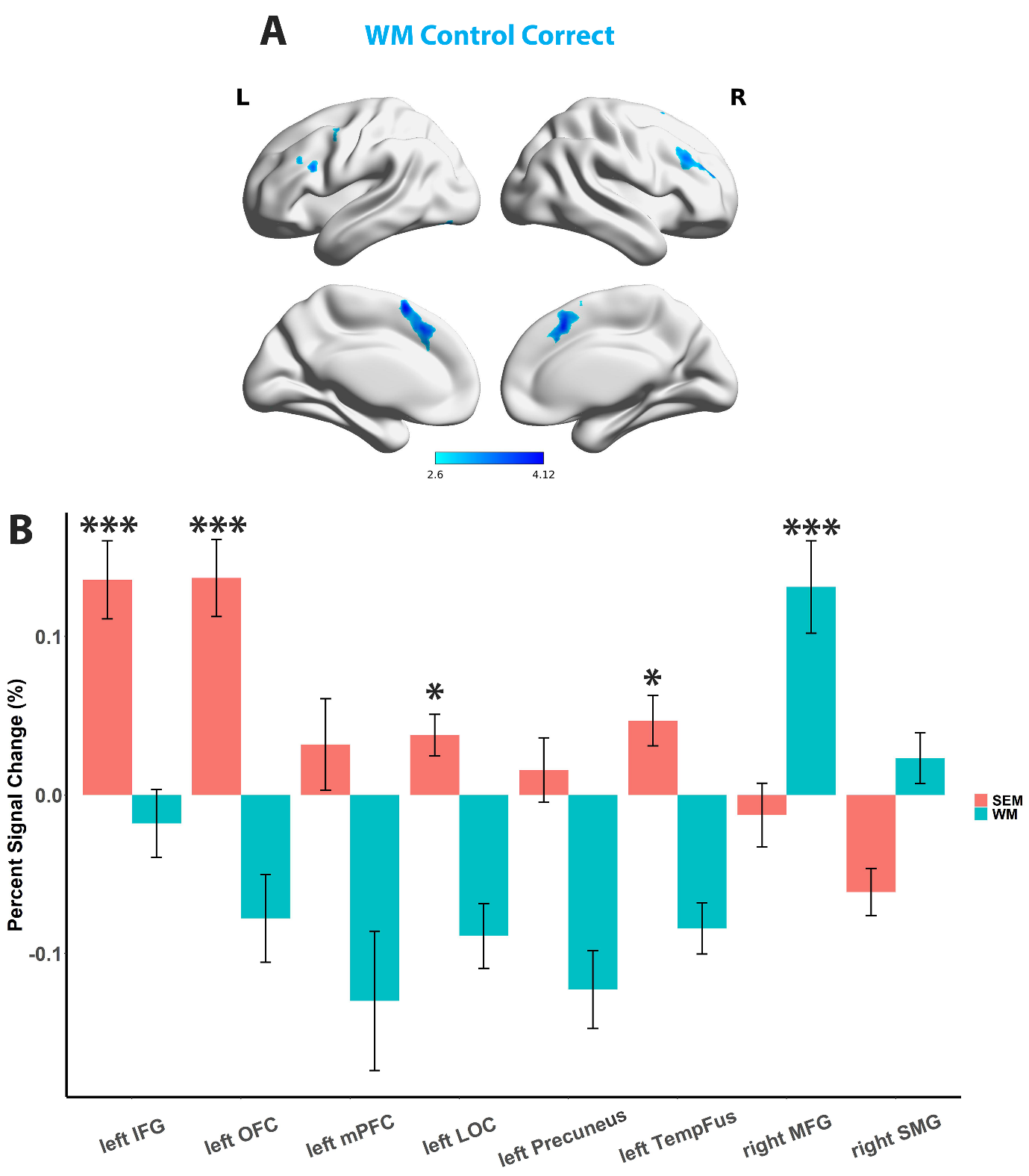

**Figure S2.** A. Parametric modulation effect of working memory load for the correct trials using a cluster detection statistics threshold of Z > 2.6. B. Left insula showed a significant positive parametric effect of semantic task difficulty, while right insula was positively modulated by working memory load. C. The clusters showing significant differences in the parametric effect of task difficulty in the main text Figure 2E and Figure 2G: inferior frontal gyrus (IFG), frontal orbital cortex (FOC), medial prefrontal cortex (mPFC), precuneus and temporal fusiform (TempFus) in the left hemisphere showed significant positive parametric modulation effect of semantic task difficulty; right middle frontal gyrus (MFG) showed significant positive parametric effect of working memory load. Bonferroni correction was used, ***p < 0.001/8. **p < 0.01/8. *p < 0.05/8.

**Neural Coding of Semantic and Working Memory Demand in Large-scale Brain Networks**

In order to test whether the same neural code supported semantic control demands and working memory load, we examined classification of control demands in each task, and cross-classification of difficulty across tasks, within MDN and SCN. We performed a series of SVR decoding analyses in ROIs selected to fall within the following networks: (i) sites within the semantic network but not implicated in control; (ii) SCN (defined as voxels within the semantic control network identified by Noonan et al. (2013) and updated by Jackson (2020), and yet outside the MDN); (iii) regions common to both SCN and MDN; (iv) MDN (defined as voxels within the multiple-demand network identified by Fedorenko et al. (2013), and not within the SCN. For the semantic task, word2vec (as a measure of relatedness) and difficulty (as assessed by behavioural performance) show an opposite relationship for trials judged to be related and unrelated. The information coded in the neural activation patterns is high-dimensional and complex, we therefore reasoned that a classifier sensitive to semantic control and/or executive demands would show a positive correlation between actual and predicted control demands when trained on related trials (which had low word2vec values for more difficult trials) and then tested on unrelated trials (which had high word2vec values for more difficult trials), or vice versa. In contrast, brain regions showing a negative correlation across these trial types would be sensitive to the associative strength of the presented pairs of items, irrespective of the subsequent judgement. Moreover, brain regions able to cross-classify difficulty between semantic and working memory tasks should be sensitive to domain-general control demands. In contrast, brain regions sensitive to difficulty in one or both tasks that do not show this task cross-classification may reflect task-specific control processes, and not domain-general control.

The within-task and between-task classification results are summarised in Figure S3. We found the difficulty of both tasks could be decoded across all four networks (compared to chance level, all p values < 0.001; all networks survived Bonferroni correction). There was also significant generalization of difficulty within the semantic domain (between semantically-related and unrelated trials) in all control networks (all p < 0.001), and also in non-control-semantic regions (p = 0.002). All p values were adjusted by Bonferroni correction.

However, this global sensitivity to difficulty across tasks and networks does not test whether the neural representation of difficulty was equivalent across tasks. Cross-classification between the semantic and working memory tasks was found only for the MDN-only regions and MDN+SCN regions for Semantic2WM (p = 0.018 and p = 0.012, respectively, which both with Bonferroni correction applied). The SCN-only and non-control-semantic regions were unable to cross-classify difficulty across these different language tasks (p = 0.594 and p = 0.891, respectively, with Bonferroni correction applied). These results indicate that semantic demand and working memory load did not share a common neural code in semantic-only networks, although they did in multiple-demand regions. This result demonstrates that semantic demand is not analogous to other types of control within the SCN.

A series of one-way repeated-measures ANOVA including the within-participant factors of brain network (with 4 levels) were used to examine the neural representation of cognitive demands for each classification type. We found a significant main effect of brain network for semantic difficulty decoding for the trials judged to be related (Related) (F(2.162,54.042) = 5.458, p = 0.006, Greenhouse-Geisser corrected). Post-hoc paired sample t-tests revealed significant better neural coding of semantic difficulty in SCN+MDN regions than non-control-semantic regions and MDN-only regions (p = 0.014 and p = 0.014 with Bonferroni correction applied, respectively), suggesting a more important role of SCN related regions than MDN regions in representing semantic control demands. There was no reliable difference of brain networks for the decoding accuracy of semantic difficulty for the trials judged to be unrelated (Unrelated) (F(2.403,60.078) = 0.429, p = 0.689, Greenhouse-Geisser corrected) and for WM load (F(2.302,57.553) = 2.363, p = 0.096, Greenhouse-Geisser corrected). There was a significant main effect of brain network for cross-condition decoding accuracy within the semantic domain (between related and unrelated decisions; F(2.305,57.623) = 5.859, p = 0.003, Greenhouse-Geisser corrected): post-hoc t-tests revealed there were reliable differences between non-control-semantic regions compared with SCN+MDN regions and MDN-only regions (p = 0.001 and p = 0.023 after Bonferroni correction, respectively), with the control regions showing better decoding. For the cross-task classification, there was a significant main effect of brain networks (F(2.462,61.551) = 6.194, p = 0.002, Greenhouse-Geisser corrected), post-hoc t-tests revealed higher decoding accuracy in SCN+MDN and MDN-only than in non-control-semantic regions (p = 0.002 and p = 0.038 with Bonferroni correction applied, respectively).

Recent studies have suggested that there is graded functional change from DMN through SCN regions to MDN: these networks form an orderly sequence on the cortical surface that is captured by the “principal gradient” of intrinsic connectivity (Margulies et al. 2016; Wang, Margulies, et al. 2020). The principal gradient has been shown to relate to the order of large-scale canonical networks on the cortical surface, from DMN, through MDN regions (frontoparietal and attention networks) to primary sensory and motor cortex. On this spectrum, the SCN falls between DMN and MDN (Wang et al., 2020). Given these previous findings, we would expect a linear change across brain networks in cross-task decoding accuracy for semantically-relevant DMN, SCN-only and SCN+MDN regions. The MDN-only regions were removed from this analysis, since this network by definition fell outside semantically-relevant cortex, and consequently included many regions that were not adjacent to SCN+MDN on the cortical surface (see Figure S3). ANOVA revealed a significant linear contrast effect in cross-task decoding accuracy across semantic not control, SCN-only and SCN+MDN regions (F(1,25)=17.032, p < 0.001). Simple t-tests revealed there was significantly higher decoding accuracy in SCN+MDN regions than semantic not control regions (p < 0.001, Bonferroni corrected), and higher cross-task decoding accuracy in SCN+MDN than SCN-only regions before multiple comparison correction (p = 0.021). The linear contrast effects were also significant in the decoding of difficulty for related semantic trials (F(1,25)=11.566, p = 0.002) and the cross-decoding between related and unrelated trials (F(1,25)=19.06, p < 0.001), and approached significance for the decoding of working memory load (F(1,25)= 3.881, p = 0.06). Thus, our results suggest graded changes in the representation of control demands from DMN through SCN to SCN+MDN. These results demonstrate the critical role of SCN regions in representing the unique neural coding of semantic control demands, alongside the roles of MDN-only and MDN+SCN regions in representing the neural coding of both semantic and non-semantic control demands.

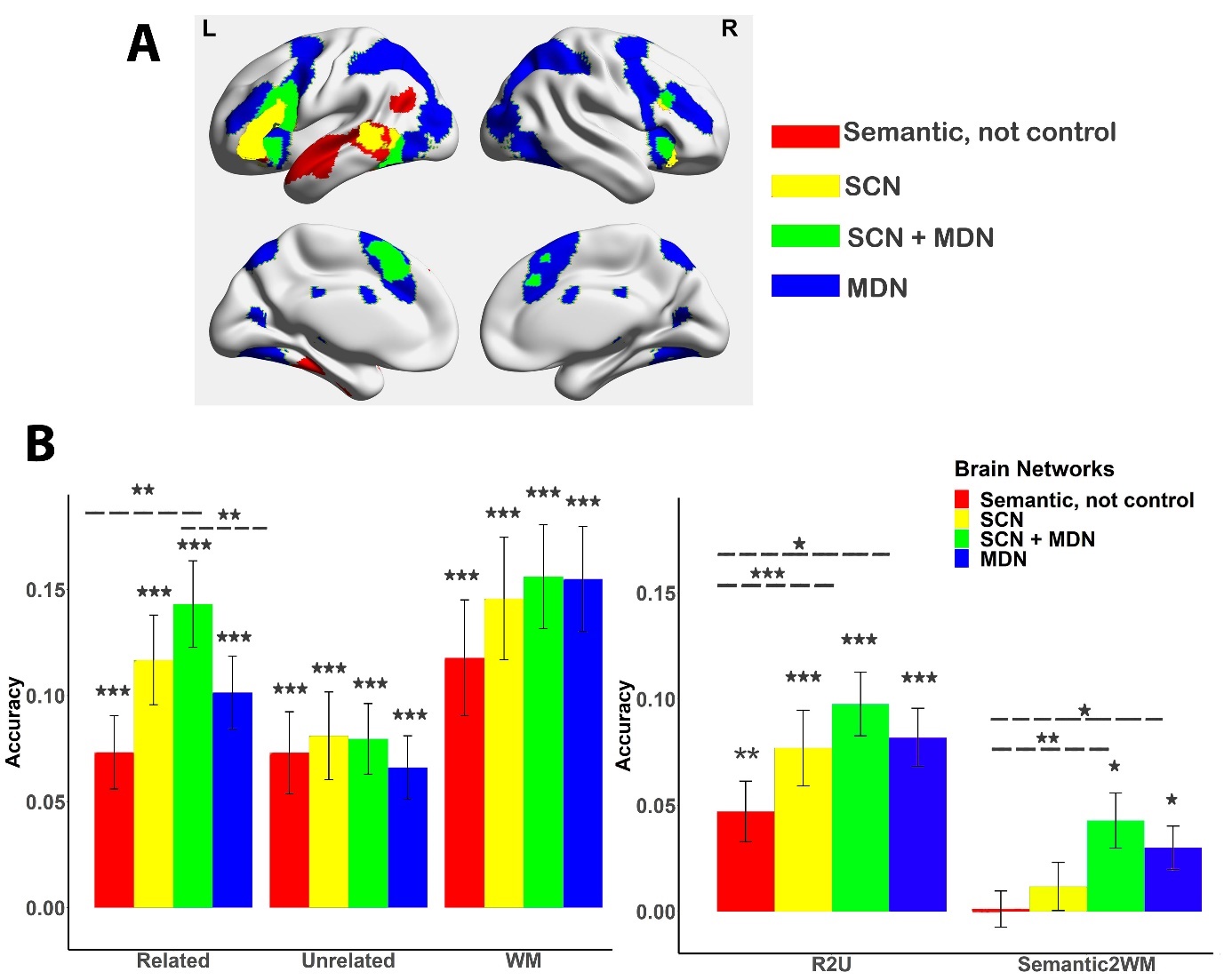

**Figure S3.** Decoding cognitive demand in large-scale brain networks; bars reflect average accuracy which is different from the conventional SVM classification and instead measures correlation (Fisher’s z-transformed) between actual task demand (difficulty) and predicted task demand. A. Brain regions for each network. B. Related: The decoding accuracy for semantically related trials. Unrelated: The decoding accuracy for semantically unrelated trials. WM: Decoding accuracy for WM correct trials. R2U: Cross-condition classification between semantic related and unrelated trials. Semantic2WM: Cross-task classification between semantic difficulty and WM load. Cross-task decoding was not significantly greater than chance level (0) in the ‘semantic, not control’ and SCN networks, and significantly higher than chance level in MDN+SCN regions and MDN (p = 0.012 and p = 0.018, respectively, with Bonferroni correction applied). All other decoding accuracy results are significantly higher than chance level in all networks (P < 0.001). Bonferroni correction was applied for each condition, separately; ***p < 0.001/4. **p < 0.01/4. *p < 0.05/4.

**Neural Coding of Semantic Demand and Working Memory Load in Large-scale Brain Networks *Randomly Choosing Voxels from Each Network***

Next, in order to confirm that our results were not underpinned by specific ROIs in each network but reflected the characteristics of neural coding of cognitive demands across the network, we randomly sampled 200 voxels in each network across ROIs in each iteration (see Figure S4).

We found difficulty could still be decoded in both semantic and WM tasks in all four networks (compared to chance level: 0, all Ps < 0.0001). There was significant generalization of difficulty within the semantic domain (between semantically-related and unrelated trials) in all four networks (all Ps < 0.0002, with Bonferroni correction applied), suggesting SCN and MDN are both sensitive to the difficulty of semantic decisions, as opposed to semantic similarity per se.

There was significant generalization across the semantic and WM tasks in MDN+SCN regions (p = 0.0016, with Bonferroni correction applied). We did not find significant cross-task classification in other networks (Semantic, not control regions: p = 0.441; SCN: p = 0.09; MDN regions: p = 0.414, with Bonferroni correction applied). This inability to cross-classify within the semantic network indicates that semantic demand and non-semantic demand do not share a common neural code.

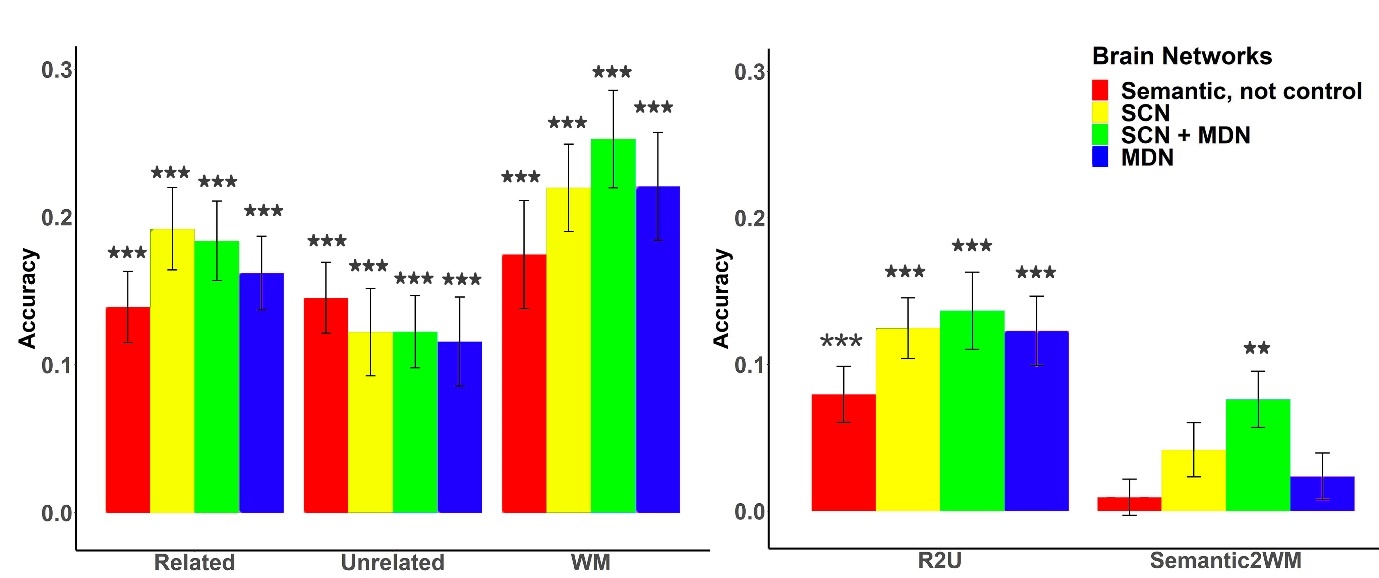

**Figure S4.** Decoding cognitive demand information in large-scale brain networks. Cross-task decoding for Semantic2WM was not significantly greater than chance level (0) in the ‘semantic, not control’ SCN networks, while significantly higher than chance level in shared (MDN+SCN) regions (p = 0.0016, with Bonferroni correction applied); there was a trend toward significant higher than chance level in MDN specific regions (p = 0.09, with Bonferroni correction applied). All other decoding accuracy results are significantly higher than chance level in all networks (all Ps < 0.0002, with Bonferroni correction applied). Bonferroni correction was used, ***p < 0.001/4. **p < 0.01/4. *p < 0.05/4.

|  | tSNR |
| --- | --- |
| LIFG | 191.867 |
| LMFG | 246.706 |
| LSFG | 277.683 |
| L anterior Cingulate Gyrus | 239.94 |
| L Insular | 215.159 |
| L anterior ITG | 144.772 |
| L Posterior Temporal gyrus | 277.246 |
| L Temporal Pole | 107.766 |
| LAG | 312.473 |
| LPrecuneous | 271.327 |

**Figure S5.** tSNR values in example ROIs. tSNR was calculated for each participant by dividing the mean of smoothed time series in each voxel by its standard deviation in each run and averaging the tSNR across all runs for the semantic task. The tSNR values were comparable with previous studies (Hoffman et al. 2015; Striem-Amit et al. 2018), and were at acceptable level (Murphy et al. 2007), although lowest at the anterior temporal pole (mean value: 107.8). LIFG: left inferior frontal gyrus; LMFG, left middle frontal gyrus; LSFG, left superior frontal gyrus; LAG, left angular gyrus.
